# Effects of temperature on the development of *Heliconius erato* butterflies

**DOI:** 10.1101/2022.12.07.519472

**Authors:** Yuqian Huang, Josie McPherson, Chris D. Jiggins, Gabriela Montejo-Kovacevich

## Abstract

1. Anthropogenic climate change is thought to present a significant threat to biodiversity, in particular to tropical ectotherms, and the effects of long-term developmental heat stress on this group have received relatively little research attention.
2. Here we study the effects of experimentally raising developmental temperatures in a tropical butterfly. We measured survival, development time, adult body mass, and wing size of a neotropical butterfly, *Heliconius erato demophoon*, across three temperature treatments.
3. Egg survival was lower in the hotter treatments, with 83%, 73%, and 49% of eggs eclosing in the 20-30°C, 23-33°C, and 26-36°C treatments, respectively. Larval survival was five times lower in the 26-36°C treatment (4%) compared to the 20-30°C treatment (22%), and we did not detect differences in pupal survival across treatments due to high mortality in earlier stages.
4. Adults in the 20-30°C treatment had a lower body mass and larvae had a lower growth rate compared to the intermediate 23-33°C treatment, but were heavier than the few surviving adults in the 26-36°C treatment. Females were heavier and grew faster as larvae than males in the 23-33°C treatment, but there was no associated increase in wing size.
5. In summary, high developmental temperatures are particularly lethal for eggs and less so for larvae, and also affect adult morphology. This highlights the importance of understanding the effects of temperature variation across ontogeny in tropical ectotherms.

## Introduction

Environmental temperature is one of the most critical ecological parameters for ectothermic species, as they have limited ability to adjust their body temperature (Sunday *et al*., 2011). Thus, both prevailing weather and long-term climate change directly impact insects, and their capacity to cope with varying temperatures is critical for species survival and dispersal. Exposure to a high temperature typically reduces individual fitness and ultimately causes death through protein denaturation, disruption of membrane structure, and desiccation (Klockmann *et al*., 2017b). Furthermore, heat stress may lead to long-term life-history trade-offs in those survivors, restricting future reproductive success (Jourdan *et al*., 2019). Under current global warming projections, high temperatures and extreme climate events will be more frequently encountered in the future, which may strongly impact biodiversity (IPCC 2021).

Although half of known animal species are tropical ectotherms, our knowledge about how well they can withstand high temperatures remains limited (García-Robledo *et al*., 2016; Sheldon, 2019). There are many studies on the effects of heat stress on the development of animals (Piyaphongkul *et al*., 2012), but fewer have focused on tropical insects. Furthermore, the majority of these studies have focussed on adults, thus, much less is known about the impact of thermal stress throughout development (Klockmann *et al*., 2017b). Tropical insects are predicted to be especially vulnerable to elevated temperatures (Janzen, 1967). Since environmental temperatures in the tropics are largely stable throughout the year, this could potentially lead to narrow thermal tolerance in tropical species (Polato *et al*., 2018; Sheldon, 2019). As such, tropical ectotherms are likely to be living near their upper-temperature limits already and may not be able to cope with a large increase in developmental temperatures (Deutsch *et al*., 2008; Fischer *et al*., 2014). Thus, it is important to determine the ability of tropical ectotherms to overcome the physiological constraints posed by increasing temperatures (Piyaphongkul *et al*., 2012).

Thermal stress experienced by juveniles may further affect adults (“carry-over effects”; (Klockmann *et al*., 2017b). Juveniles surviving heat stress may have reduced fitness as adults, and could influence life-history trade-offs (Jourdan *et al*., 2019). Under elevated temperatures, there may be a trade-off in phenology and morphology, resulting in shorter development time and emergence at a lower adult body size (Chown & Gaston, 2010; Fischer *et al*., 2014). Adult body size is often a predictor of lifespan, competitiveness, and reproductive success, possibly impacting population viability (Fischer *et al*., 2004; Kingsolver & Huey, 2008). Therefore, carry-over effects from juvenile heat stress could severely impact adult fitness. Improving our understanding of tropical insect responses to global warming will necessitate considering the effects of heat stress on both survival and other fitness traits throughout development (Klockmann *et al*., 2017b).

This study investigates the effects of increased temperatures on survival and development in a Neotropical butterfly *Heliconius erato demophoon. H. erato* is a pollen-feeding butterfly ranging from southern Texas to northern Paraguay and often inhabits fringes of tropical rainforests (Turner, 1971). Previous studies have shown plasticity in heat tolerance when butterflies from different elevations were reared in a common-garden environment, with butterflies from different populations showing similar heat tolerances despite strong differences in the wild (Montejo-Kovacevich *et al*., 2020). Here we test the vulnerability of each juvenile stage to sustained heat stress by testing the survival and growth rate of eggs, larvae, and pupae. We then investigate the effects of elevated temperatures on other traits in the surviving adults, including adult mass and wing development.

## Materials and Methods

### Study system

We used an established laboratory stock population at the Madingley rearing facility, University of Cambridge, UK, that was started in 2017 with *Heliconius erato demophoon* butterflies from Panama. The larvae of the stock population were reared at 25°C, 75% relative humidity, and a photoperiod of L12:D12 within a single temperature-, light-, and humidity-controlled CT room. The adults were kept in the insectaries, which experienced variable diurnal temperature changes. The larvae were fed with tips of *Passiflora biflora*, and adults were fed with pollen from *Lantana camara* and supplemented with protein-incorporated sugar solution.

### Experimental design

Eggs were collected from dozens of females in the stock population in the summer of 2021. Since the stock populations had been reared and bred together for multiple generations, effects due to genetic differences between parents were considered negligible, and thus, we did not track pedigrees in this experiment. However, each egg was tracked individually through development.

The host plants *P. biflora* were removed from the cages with adults 24-48 hours before collecting eggs, to encourage females to lay when the host plants were reintroduced. Approximately 20 *H. e. demophoon* females were allowed to oviposit on a small *P. biflora* plant for a three-hour session on three consecutive days, from which the eggs were collected hourly, and time was noted. Three eggs were placed per small plastic pot. Pots were randomly placed among three treatment groups within 2 hours of laying (Figure 1). Climate chambers were heated up to the target temperature before the transfer of pots, and the light: dark, and temperature cycles were 12h:12h across cabinets. The 20-30°C temperature (i.e., 12h of 20°C followed by 12h of 30°C, also referred to as the cold treatment) was used as a control for the comparison, with the same mean temperature as the normal rearing temperature of the stock population, i.e. 25°C. The 23-33°C temperature (also referred to as the intermediate treatment) used is within the range of temperature experienced by *H. e. demophoon* in some locations of its natural habitat, and 26-36°C (also referred to as the hot treatment) represents temperatures only experienced occasionally in forest canopies in the Equator (Montejo-Kovacevich *et al*., 2020) but that could become common in degraded lowland habitats in the tropics (Luber & McGeehin, 2008).

**Figure 1.**
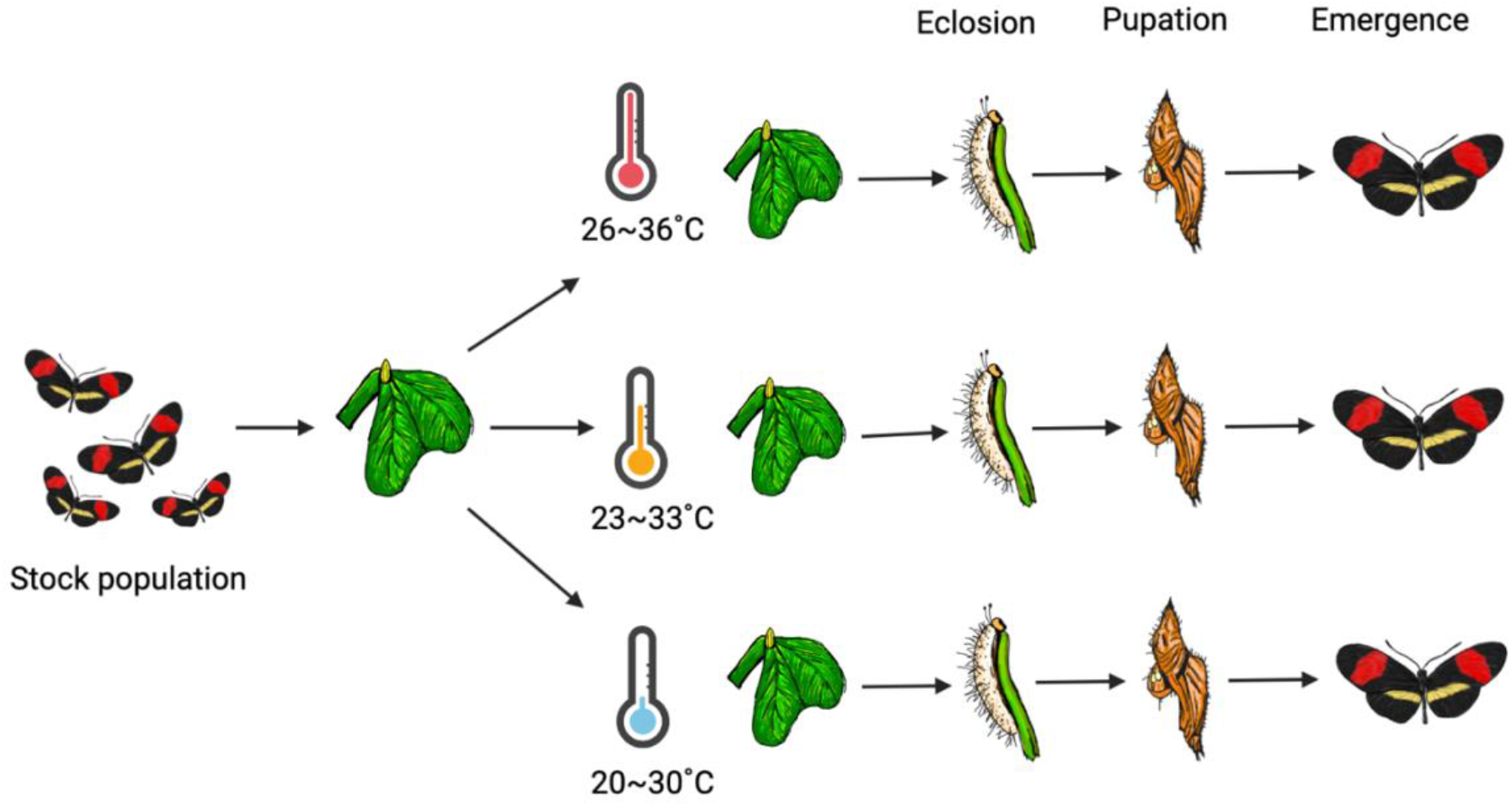
Experimental design. The stock population was allowed to oviposit on the host plant, *Passiflora biflora*, and the eggs were randomly separated into three temperature treatment groups (20-30°C, 23-33°C, or 26-36°C). The time of eclosion, pupation, and emergence were tracked, as well as individual survival and mass.

Survival rates and development times for eggs, larvae, and pupae were measured (Figure 1). Egg eclosions were checked three times a day at 10 am, 1 pm, and 4 pm for seven days after laying and time was noted. Eggs were considered dead if no larvae had emerged after seven days. The larvae from each temperature group were transferred into plastic boxes lined with moist tissue and *P. biflora* with three to five larvae per box and survival was checked every two days. Pupation and emergence dates were recorded to obtain larval and pupal development time. Pupae were weighed between 2 and 4 days after pupation. Each pupa was hung in an individual, labelled pot once weighed. After emergence, butterfly wings were carefully removed from the thorax with forceps to best maintain the whole structure. Adult body mass was measured after wing removal, and bodies were stored in pure ethanol at −20°C for future studies.

### Wing image analysis

Wing size was obtained from images of the detached wings of the individuals that reached adulthood (n=78). Detached wings were photographed dorsally and ventrally with a DSLR camera with a 100 mm macro lens in standardised conditions. Any damaged or folded specimens were excluded from the analysis. A custom script for Fiji (Schindelin *et al*., 2012), which automatically crops, extracts the forewings, and performs particle size analysis, was used to obtain wing measurements from the images (Montejo-Kovacevich *et al*., 2019a). We obtained an average wing area between the forewings (where possible, in mm^2^, hereafter “size”) since butterflies predominantly use their forewings for flight (Le Roy *et al*., 2019).

### Statistical analysis

We analysed (1) the survival rates of eggs/larvae/pupae as the percentage of eclosion/pupation/emergence per treatment group, respectively, (2) development time across treatments, (3) pupal mass and adult body mass per treatment group and sexes, and (4) wing size and wing loading between treatment groups and sexes. Pairwise t-tests were performed for the means of each treatment group. Bonferroni adjustments were performed after chi-squared tests if the group number was more than two. All statistical analyses were run in R V1.3.1093 (R Development Core Team 2011), and graphics were generated with the package ggplot2 (Ginestet, 2011).

## Results

### Survival and development time

The survival rate of all stages decreased under heat stress (Figure 2). Egg and larval survival were both affected by heat stress (X^2^ (2, *N* = 928) = 74.95, *p* < 0.0001, and X^2^ (2, *N* = 633) = 16.01, *p* < 0.001, respectively). Post hoc pairwise comparisons of survival revealed that egg survival was lower in the two hotter treatments (egg survival was 83.0%, 73.3%, and 48.6%, respectively) whereas larval survival was only significantly lower in the 26-36°C treatment compared to the 20-30°C treatment (larval survival was 22.2%, 14.0%, and 3.8%, respectively). We did not detect differences in pupal survival across treatments, but sample sizes were lower due to mortality in earlier stages (X^2^(2, *N* = 144) = 4.40, *p* = 0.11,).

**Figure 2.**
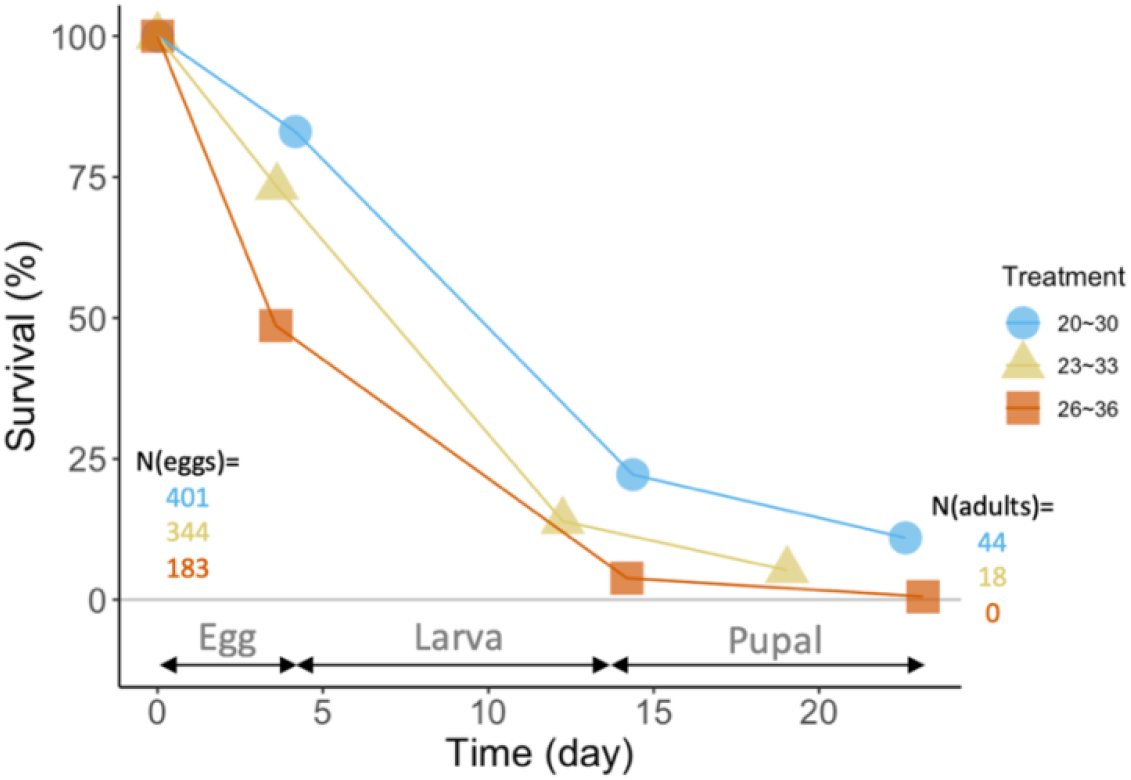
Percentage of individuals surviving during egg, larval, and pupal stages in three temperature treatments (blue = 20-30°C, yellow = 23-33°C, and red = 26-36°C).

Timing of larval death was not correlated with rearing temperature. However, individuals that died as pupae in the 20-30°C treatment took around 30 hours longer to develop as larvae compared to those that successfully emerged as adults in the same treatment (t(41) = 2.55, *p* = 0.014, and t(53) = 2.87, *p* = 0.0059 larval development time of females/males that reached adulthood compared to that of those that died as pupae, classed as unknown sex in Fig 3A treatment 20-30°C, respectively).

**Figure 3.**
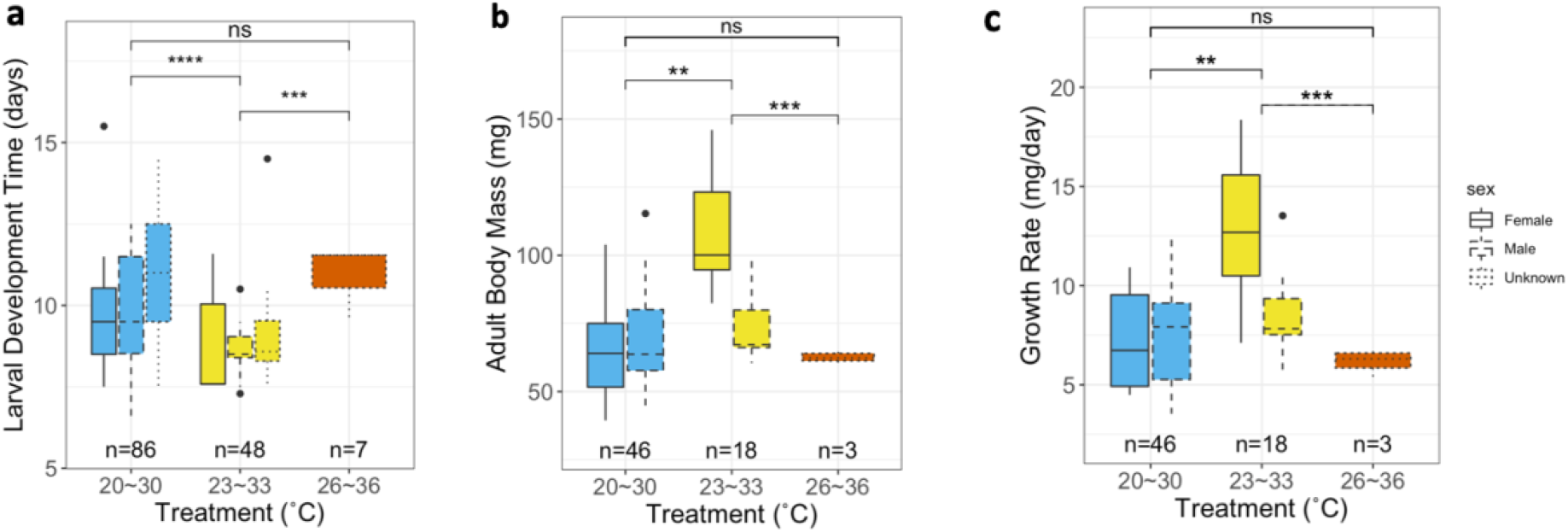
Growth and development across treatments and sexes: (a) larval development time, (b) adult body mass (mg), and (c) growth rate measured as mass gain (mg) per day. Blue, yellow and orange bars represent 20-30°C, 23-33°C and 26-36°C treatment, respectively. Solid lines are females and dashed lines are males. Top values showing significant levels of pairwise t-tests between each treatment group (o < 0.1, * < 0.05, ** < 0.01, *** < 0.001). Individuals that died at the pupal stage were not sexed (‘Unknown’ sex category).

Overall development time was fastest in the intermediate treatment (19.0 ± 1.4 days) and similar in the cold and hot treatments (22.6 ± 1.9 days and 23.1 ± 0.6 days, respectively). There was no significant difference in development times between sex in each treatment group (p = 0.70).

### Mass and growth rate

Adults were heaviest in the intermediate treatment (for 20-30°C and 23-33°C, t(24) = 2.87, p < 0.01; 23-33°C and 26-36°C, t(18) = 4.04, p < 0.001), but the cold and hot treatments yielded adults with similar body mass (t(21) = 1.89, p = 0.07, Figure 3B). Nevertheless, when comparing adults of the same sex between treatments, females in the intermediate treatment were heavier than females in the cold treatment (t(8) = 4.21, p < 0.01), whilst males in the cold and intermediate groups did not significantly differ in weight. Further, females in the intermediate group were significantly heavier than males in the same treatment (t-test between sexes in 23-33°C, t(7) = 3.70, p < 0.01). Thus, the difference in mean body mass between cold and intermediate treatment was driven by heavier females in the intermediate temperatures. Pupal weight was similar between the cold and intermediate groups but decreased significantly at 26-36°C (for 20-30°C and 26-36°C, t(16) = 3.06, p < 0.01, and 23-33°C and 26-36°C, t(20) = 4.28, p < 0.001).

Growth rate, which is the ratio between adult body mass and larval development time (since growth only occurs at the larval stage), was highest at 23-33°C and decreased at 26-36°C compared to the 20-30°C treatment (for 20-30°C and 23-33°C, t(22) = 3.21, p < 0.01, and 23-33°C and 26-36°C, t(17) = 4.26, p < 0.001, Figure 3C). However, only females from the 23-33°C had an elevated growth rate compared to the females from the 20-30°C treatment (t(17) = 4.26, p < 0.001), not the males. Since there was no difference between development times, females in higher temperatures grew both faster and heavier. In summary, a small increase in temperature led to a faster growth rate, but the growth rate dropped when the temperature was too high.

### Wing size and loading

There was no difference in wing size between temperature treatments or sexes (R^2^_LM_ = −0.02, F(2, 59) = 0.42, p = 0.66; Figure 4). The hot treatment was excluded from the comparison as most individuals in this treatment did not reach adulthood. Temperature treatment and sex were both predictors of wing loading, which is the ratio between the adult body mass and wing size (R^2^_LM_ = 0.27, F(2, 57) = 12.15, p < 0.0001). Females in the 23-33°C treatment had a greater wing loading than males in the same group (t(5) = 2.78, *p* = 0.04). These differences in wing loading were likely driven by the higher adult body mass in the warmer treatment since there was no difference in wing size.

**Figure 4.**
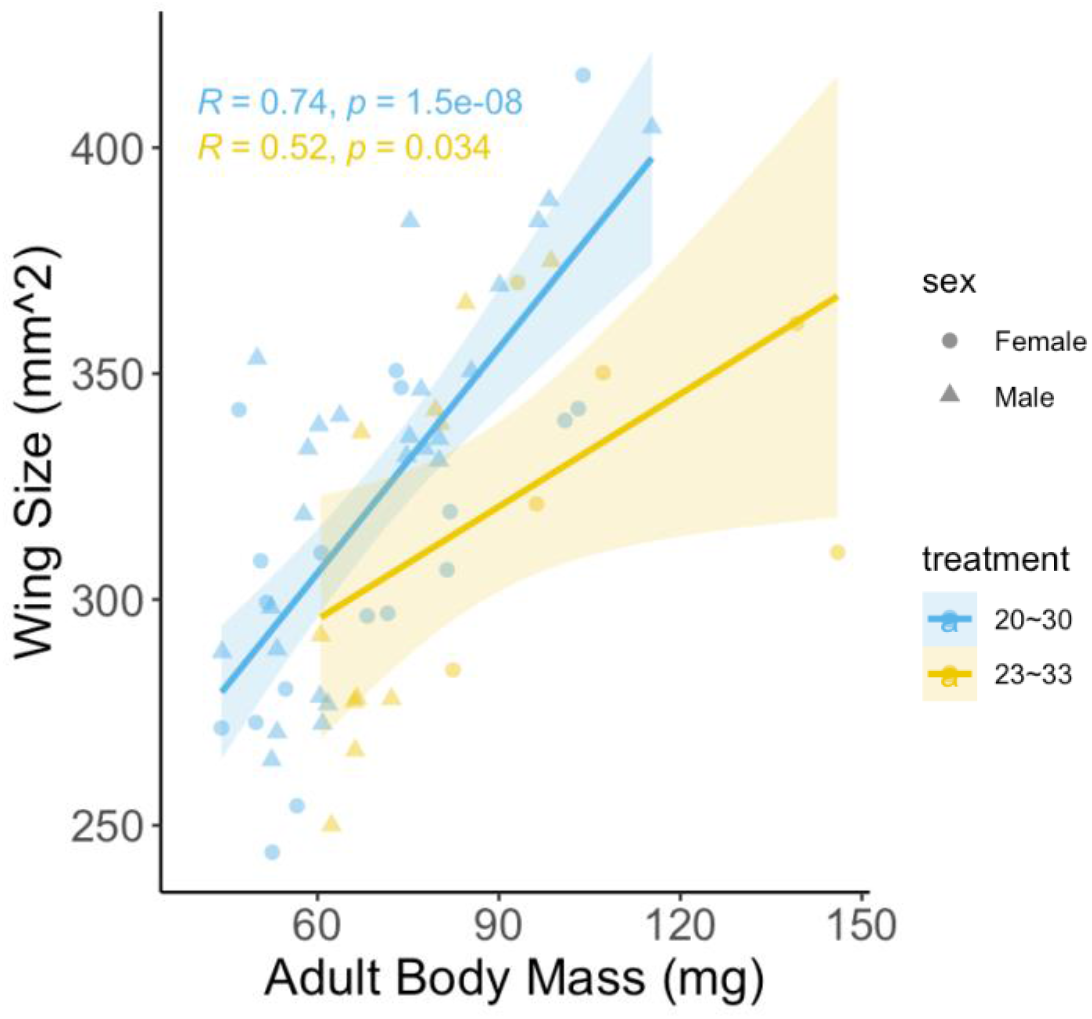
Adult wing size and body mass allometry. Wing analyses between treatment groups (blue = 20-30°C, yellow = 23-33°C). Pearson correlation coefficients and p-values are shown for the regression lines per treatment.

## Discussion

Insects have developed a variety of adaptations to endure temperature variation (Overgaard *et al*., 2008), yet there are temperature thresholds beyond which species cannot live. This study revealed that a 3°C temperature increase to 23-33°C led to higher egg mortality and a 6°C temperature increase to 26-36°C led to both higher egg and larval mortality compared to developing at 20-30°C. The few survivors of 26-36°C had a significant decline in adult body mass, growth rate, and wing loading. In contrast, the slight elevation from 20-30°C to 23-33°C increased adult body mass and growth rate, but the fitness consequences of these trait shifts were not assessed.

### Temperature affects survival and development

In line with earlier studies, we found strong adverse effects of high temperature on both egg-hatching success and larval survival (Piyaphongkul *et al*., 2012; Kingsolver *et al*., 2015; Klockmann *et al*., 2017a). These detrimental effects could be explained by, for instance, denaturation of proteins, disruption of membrane structure, or desiccation (Klose & Robertson, 2004; Chown & Terblanche, 2006; Potter *et al*., 2009). The *H. e. demophoon* larvae here studied seems to be well equipped to bear temperatures slightly above the normal rearing temperature (25 °C) of the stock, as there were no differences in larval and pupal mortality between the 20-30°C and 23-33°C treatments.

Furthermore, it is remarkable that some individuals survived cycles of 12 hours at 36°C in the 26-36C treatment, as individuals of the same species have been shown to get knocked down after only 15.9 minutes when exposed to temperatures between 39 and 41°C (Montejo-Kovacevich *et al*., 2020). Many studies have looked at the maximum temperature organisms can endure (critical thermal maximum) or perform short exposures to high temperatures and measured the time until they are knocked down (Bowler & Terblanche, 2008; Ju *et al*., 2014; Nandi & Chakraborty, 2015). Our findings suggest that responses to short exposures may not correlate so well to long exposures. In the wild, short exposures may be relevant to complex habitats such as tropical forests where microclimates are buffering against heat stress, acting as refugia (Scheffers *et al*., 2014). But for other more open or degraded habitats, temperature increases may be more homogenous throughout space and time of the day (De Frenne *et al*., 2019; Montejo-Kovacevich *et al*., 2020). Therefore, testing different exposure times and temperatures for different habitat types would be interesting for future studies.

The ability to survive in hot conditions changes through development, with eggs having the lowest survival, followed by larvae. Different hypotheses as to why heat tolerance varies through development have been proposed. First, individuals with a larger size may have higher survival under heat stress (Kingsolver & Huey, 2008; Klockmann *et al*., 2017a). Klockmann *et al*. (2017b), for example, found that survival was higher for larger caterpillars of *Bicyclus anynana*. Nonetheless, we did not find size-dependency in heat tolerance within larval development as the time of larval death did not differ between the treatment groups. Second, the enhanced survival of larvae compared to eggs may be attributed to behavioural responsiveness. Since desiccation is an important factor contributing to the high mortality under heat stress, the consumption of food by the larvae may allow them to acquire water and energy which are essential for evaporative cooling to prevent desiccation (Klockmann *et al*., 2017a). Finally, habitat use by wild butterflies may further explain why eggs are sensitive to high temperatures. Eggs may not be under strong selection pressure to evolve heat tolerance in the wild if they inhabit the understory and near the moist leaves, which can provide microclimate buffering against heating (Madigosky, 2004). The understory of tropical forests inhabited by *Heliconius* in Ecuador has, on average, 1.75°C cooler daily maximum temperature than the sub-canopy (Montejo-Kovacevich *et al*., 2020). The absence of such refugia in our experiment, with only a few leaves and/or stems of the plant in each pot, may have exacerbated the heat stress suffered by the eggs. Overall, our findings are broadly in line with similar experiments on *Bicyclus anynana* butterflies and highlight the importance of preserving habitat complexity for buffering against heat stress in vulnerable and immobile early life stages of tropical ectotherms.

Our results showed that adult body mass was higher at 23-33°C relative to the 20-30°C treatment, but lower at 26-36 °C. Since temperature is known to have a major influence on various developmental processes in ectotherms (Lailvaux & Irschick, 2007), higher temperatures may lead to butterflies reaching adulthood faster. In line with previous studies, the overall trend of growth rate is similar to a typical temperature-performance curve, with higher growth rates in intermediate to low temperatures (Huey & Stevenson, 1979; Lee & Roh, 2010).

Growth is defined as the net energy surplus of absorption and metabolism, and thus, the growth rate could be modelled by the relationship between a series of processes, including metabolism and ingestion (Wood & McDonald, 1996). As temperature increases, larvae may increase their ingestion rate to compensate for the increased metabolic cost (Kingsolver *et al*., 2015). The initial increase in ingestion rate may over-compensate the increase in metabolic cost if the food supply is not a limiting factor, giving rise to the boosted growth rate and fitness benefits seen at 23-33°C. However, this ingestion rate will eventually reach a plateau and decline beyond the optimum temperature, which, in this study, is likely to be around 28°C (the mean of 23-33°C) for *H. e. demophoon*. The reasons for the decline in ingestion could be those similar reasons involved in the increased mortality rate under heat stress, eventually leading to smaller adult body mass under high developmental temperatures. Whilst growth rate results match this expected metabolism-ingestion relationship, we did not explicitly measure the rate of metabolism and ingestion for individual caterpillars across the temperature treatments. Since the larvae in our experiments were allowed to feed *ad libitum*, we do not know exactly how the rate of ingestion and metabolism are affected by temperature. Future studies may look into the difference in the amount of feeding and the resulting adult body mass between temperature treatments for empirical validation of the metabolism-ingestion relationship.

### Females respond more plastically to temperature

Our results found that under mildly elevated temperatures (23-33°C), females had a greater increase in body mass and growth rate than males (Figures 3B and 3C). Females often have a higher body mass due to a positive correlation between body size and fecundity, whereas males are typically selected for larger wings, which increases their mating opportunities (Deinert *et al*., 1994; Gotthard *et al*., 1994; Blanckenhorn *et al*., 2007; Montejo-Kovacevich *et al*., 2019b). This can be explained by the fact that as female reproductive success is more closely correlated with adult body mass, female butterflies tend to weigh more than male butterflies under fecundity selection (Blanckenhorn *et al*., 2007; Gotthard, 2008). Thus, the optimal adult body mass differs between the sexes. Furthermore, there were no developmental time differences between the sexes. This further supports the idea that females are more plastic than males under mildly elevated temperatures, leading to increased body size and growth rate (Figure 3C) (Gotthard *et al*., 1994).

Wing loading, which is the mass carried per unit area of wing size, was higher in both sexes under elevated temperatures (23-33°C). Greater wing loading may translate into reduced flight performance and dispersal ability as shown in experiments with Monarch butterflies (Soule *et al*., 2020), potentially leading to lower fitness and reproductive success despite the enhanced growth rate under warmer temperatures. Similar results of the temperature-wing loading relationship are also found in *Drosophila* (Fraimout *et al*., 2018). While adult body mass is correlated with higher egg production and fitness, it may not benefit the butterflies in natural environments if their flight ability is hindered by higher wing loading (Almbro & Kullberg, 2012). Butterflies may benefit from lower wing loading to escape predators, locate host plants, or acquire mates (Molleman *et al*., 2020). For example, higher temperature reduced host-foraging motivation in parasitoid *Aphidius colemani* (Jerbi-Elayed *et al*., 2022) and spring generation of butterflies with more significant wing loading often disperses less (Fric & Konvic, 2002; Gibbs *et al*., 2011). Male butterflies may be under stronger selection to maintain their flight ability for acquiring mates and keeping territories even when the temperature is no longer a limiting factor for body mass. Thus, heavier wing loading due to greater body mass under mildly elevated temperatures may be disadvantageous to male butterflies in the wild.

## Conclusions

This study investigated the effect of temperature on the early development of butterflies and the resulting adults, which has been rarely studied in tropical insects. Our findings reveal that long-term sub-lethal heat stress has deleterious effects on both survival and developmental traits in a common neotropical butterfly species. Eggs were found to be most vulnerable to heat stress, followed by larvae. Furthermore, the growth rate followed a typical thermal-performance curve, which showed an optimal growth temperature of 23-33°C. The observed decline in adult development rate and body mass at 26-36°C, which is known to be correlated with lower fitness (Piyaphongkul *et al*., 2012; Klockmann *et al*., 2017b), is of particular concern for two reasons. First, body mass may be a critical restriction on *H. e. demophoon* heat stress tolerance in general (Klockmann *et al*., 2017a, 2017b). Second, because smaller females often produce fewer and/or smaller eggs, this may have a transgenerational effect, resulting in offspring with lower fitness (Kingsolver & Huey, 2008; Klockmann *et al*., 2017b). The faster development rate at 23-33°C may appear to be beneficial for fecundity, but it will restrict the butterfly’s flight capabilities since wing size is less plastic than body mass. Ongoing climate change, rising ambient temperatures, and an increase in the frequency of extreme weather events such as heatwaves and drought periods would almost certainly have severe effects on life on Earth (Klockmann *et al*., 2017b; Theng *et al*., 2020). This study emphasises the importance of taking plastic responses of phenotypic traits into account when predicting population viability in response to recent global warming. It calls for incorporating the effects of temperature on developmental time and life-history traits into models forecasting species extinction risks (Jourdan *et al*., 2019).

## Supporting information

R.Script

Data set

## Acknowledgements

We would like to thank all research assistants that have contributed to this study, particularly Glennis Julian.

## Contribution of authors

Project design: Y.H., J.M., C.D.J., G.M.-K.; Data collection: Y.H., J.M., G.M.-K.; Data Analysis: Y.H., G.M.-K.; Writing – original draft: Y.H.; Writing – review and editing: Y.H., J.M., C.D.J., G.M.-K.

## Data availability

The data that supports the findings of this study are available in the supplementary material of this article.

