## Supplementary material for "Effects of temperature on the development of *Heliconius erato* butterflies": R.Script

```
library(ggplot2)
library("ggnewscale")
library(EnvStats)
library(ggbeeswarm)
library(ggpubr)
library(readxl)
library(lsr)
library("ggfortify")
library(ggpattern)


#Name dataset as Data_master
my_comparisons <- list( c("23~33", "26~36"), c("20~30", "23~33"),
c("26~36", "20~30") )
treatment<-as.factor(Data_master$treatment)


#Survival Rate plot

ggplot(data=survival_plot, aes(x=as.numeric(Time/24), y=Survival,
group=Treatment, color=Treatment))+
  geom_hline(yintercept=0, linetype="solid", color = "grey")+
  geom_point(aes(shape=Treatment),size=6, alpha=0.75)+
  geom_path(aes(col=Treatment))+
  scale_color_manual(values=c("#56B4E9","#DDCC77", "#D55E00"))+
  theme_classic()+
  ylab("Survival (%)")+
  xlab("Time (day)")+
  theme(axis.text.x=element_text(size=14),
        axis.text.y=element_text(size=14),
        axis.title=element_text(size=16))+
  geom_hline(yintercept=-10, linetype="dashed", color = "white")


#Chi squared test for Eclosion Success
chi.eclosion <- table(treatment,Data_master$eclosion.success)
chi.eclosion <- chi.eclosion[c(1:3),]
ftable(chi.eclosion)
summary(chi.eclosion)
chisq.test(chi.eclosion[c(1,2),],correct=FALSE)
chisq.test(chi.eclosion[c(1,3),],correct=FALSE)
chisq.test(chi.eclosion[c(2,3),],correct=FALSE)


chi.cat <- table(treatment,Data_master$cat.success)
chi.cat <- chi.cat[c(1:3),]
ftable(chi.cat)
summary(chi.cat)
chisq.test(chi.cat[c(1,2),])
chisq.test(chi.cat[c(1,3),])
chisq.test(chi.cat[c(2,3),])



chi.pup <- table(treatment,Data_master$pupua.success)
chi.pup <- chi.pup[c(1:3),]
ftable(chi.pup)
summary(chi.pup)
```

```
chisq.test(chi.pup[c(1,2),])
chisq.test(chi.pup[c(1,3),])
chisq.test(chi.pup[c(2,3),])



#Larva development time plot

ggplot(data=subset(Data_master, treatment %in% c("20~30", "23~33",
"26~36"),
                   pupation.eclosion.time!="" & !
(is.na(pupation.eclosion.time))),
       aes(x = treatment, y =
as.numeric(as.character(pupation.eclosion.time))/24,
       fill=treatment, linetype = sex)) +
  geom_boxplot(fill =
c("#56B4E9","#56B4E9","#56B4E9","#F0E442","#F0E442","#F0E442",
"#D55E00"),fatten=1)+
  scale_linetype_manual(values= c("Female" ="solid",
"Male"="dashed","Unknown"="dotted"))+
  #scale_alpha_manual(name = "Sex",values=c(1,0.6,0.2))+
  #geom_beeswarm(fill="grey", cex = 1.5, size=2, alpha=0.2, dodge.width=0)
+
  new_scale("linetype")+
  scale_linetype_manual(values= c("solid"))+
  stat_compare_means(method="t.test", comparisons = my_comparisons,size=5,
label="p.signif")+
  stat_n_text(size=5)+
  theme_bw()+
  ylab("Larval Development Time (days)")+
  xlab("Treatment (˚C)")+
  theme(axis.text.x=element_text(size=14),
        axis.text.y=element_text(size=14),
        axis.title=element_text(size=16))


#Larval development plot for 20-30 only
compare <- list( c("Male", "Female"), c("Male", "Unknown"), c("Female",
"Unknown") )
ggplot(data=subset(Data_master, treatment %in% c("20~30"),
                   pupation.eclosion.time!="" & !
(is.na(pupation.eclosion.time))),
       aes(x = sex, y = as.numeric(as.character(pupation.eclosion.time))/
24)) +
  geom_boxplot(fatten=1)+
  scale_linetype_manual(values= c("Female" ="solid",
"Male"="dashed","Unknown"="dotted"))+
  #scale_alpha_manual(name = "Sex",values=c(1,0.6,0.2))+
  #geom_beeswarm(fill="grey", cex = 1.5, size=2, alpha=0.2, dodge.width=0)
+
  new_scale("linetype")+
  scale_linetype_manual(values= c("solid"))+
  stat_compare_means(method="t.test", comparisons = compare ,size=5)+
  stat_n_text(size=5)+
  theme_bw()+
  ylab("Larval Development Time (days)")+
  xlab("Treatment (˚C)")+
  theme(axis.text.x=element_text(size=14),
```

```
        axis.text.y=element_text(size=14),
        axis.title=element_text(size=16))+
  stat_summary(fun = mean, geom = "text", col = "red",
               vjust = 1.5, aes(label = paste("Mean:", round(..y.., digits
= 1))))



#Adult body weight plot
ggplot(data=subset(Data_master, treatment %in% c("20~30","23~33","26~36"),
                   body.adult.mass.g!="" | !(is.na(body.adult.mass.g))),
       aes(x = treatment, y =
as.numeric(as.character(body.adult.mass.g))*1000)) +
  geom_boxplot(col=c("dodgerblue","orange", "firebrick1"))+
  geom_beeswarm(aes(colour=sex), cex = 2, size=2.75, alpha=0.5,
dodge.width=1 )+
  stat_compare_means(method = "anova", label.y = 190, label.x=0.7,size=4)+
  stat_compare_means(label="p.signif",  method="t.test", comparisons
=my_comparisons,
                     size=5,  vjust=-0)+
  stat_n_text(size=5)+
  theme_bw()+
  ylab("Adult Body Weight (mg)")+
  xlab("Treatment (°C)")+
  theme(axis.text.x=element_text(size=14),
        axis.text.y=element_text(size=14),
        axis.title=element_text(size=16))


ggplot(data=subset(Data_master, treatment %in% c("20~30","23~33",
"26~36"),
                   body.adult.mass.g!="" | !(is.na(body.adult.mass.g))),
       aes(x = treatment, y =
as.numeric(as.character(body.adult.mass.g))*1000 ,sfill=treatment,
linetype=sex)) +
  geom_boxplot(fill=c("#56B4E9","#56B4E9","#F0E442","#F0E442",
"#D55E00"),lwd=0.5, fatten=1)+
  scale_linetype_manual(values= c("Female" ="solid",
"Male"="dashed","Unknown"="dotted"))+
  stat_n_text(size=5)+
  theme_bw()+
  ylab("Adult Body Mass (mg)")+
  xlab("Treatment (°C)")+
  theme(axis.text.x=element_text(size=14),
        axis.text.y=element_text(size=14),
        axis.title=element_text(size=16))+
  new_scale("linetype")+
  scale_linetype_manual(values= c("solid"))+
  stat_compare_means(  method="t.test", comparisons =my_comparisons,
size=6,  vjust=0.5,linetype=1)+
  geom_signif(comparisons = list(c("20~30", "26~36")), annotations="ns",
              y_position = 180, tip_length = 0.03, textsize =4.5, vjust=0)


ggplot(data=subset(Data_master, treatment %in% c("23~33"),
                   body.adult.mass.g!="" | !(is.na(body.adult.mass.g))),
       aes(x = sex, y = as.numeric(as.character(body.adult.mass.g))*1000
,sfill=treatment, linetype=sex)) +
```

```
  geom_boxplot(lwd=0.5, fatten=1)+
  stat_n_text(size=5)+
  theme_bw()+
  ylab("Adult Body Mass (mg)")+
  xlab("Treatment (˚C)")+
  theme(axis.text.x=element_text(size=14),
        axis.text.y=element_text(size=14),
        axis.title=element_text(size=16))+
  new_scale("linetype")+
  scale_linetype_manual(values= c("solid"))+
  stat_compare_means(  method="t.test",size=6,  vjust=0.5,linetype=1)


#Growth Rate plot
ggplot(data=subset(Data_master, treatment %in% c("20~30","23~33",
"26~36"),
                   body.adult.mass.g!="" | !(is.na(body.adult.mass.g))),
       aes(x = treatment, y = as.numeric(body.adult.mass.g)*1000/
(as.numeric(pupation.eclosion.time)/24),sfill=treatment, linetype=sex)) +
  geom_boxplot(fill=c("#56B4E9","#56B4E9","#F0E442","#F0E442",
"#D55E00"),lwd=0.5,fatten=1)+
  #geom_beeswarm(aes(colour=sex), cex = 1, size=1.5, alpha=0.6,
dodge.width=1 )+
  scale_linetype_manual(values= c("Female" ="solid",
"Male"="dashed","Unknown"="dotted"))+
  stat_compare_means(method = "anova", label.y = 25, size=4,label.x=0.7)+
  stat_compare_means(label="p.signif",  method="t.test", comparisons
=my_comparisons,
                     size=6,  vjust=0.5)+
  #stat_compare_means(aes(group=sex), method="t.test")+
  stat_compare_means(method = "anova", label.y = 23, size=5,label.x=0.7)+
  stat_n_text(size=5)+
  theme_bw()+
  ylab("Growth Rate (mg/day)")+
  xlab("Treatment (˚C)")+
  theme(axis.text.x=element_text(size=14),
        axis.text.y=element_text(size=14),
        axis.title=element_text(size=16))+
  new_scale("linetype")+
  scale_linetype_manual(values= c("solid"))+
  stat_compare_means(label="p.signif",  method="t.test", comparisons
=my_comparisons, size=6,  vjust=0.5,linetype=1)+
  geom_signif(comparisons = list(c("20~30", "26~36")), annotations="ns",
            y_position = 23, tip_length = 0.03, textsize =4.5, vjust=0)



#Growth rate comparison by sex
sex<-as.factor(Data_master$sex)
male_only<-subset(Data_master, sex=="Male")
ggplot(data=subset(male_only, treatment %in% c("20~30","23~33"),
                   body.adult.mass.g!="" | !(is.na(body.adult.mass.g))),
       aes(x = treatment, y = as.numeric(body.adult.mass.g)*1000/
(as.numeric(pupation.eclosion.time)/24))) +
  geom_boxplot(col=c("dodgerblue","orange"))+
  geom_beeswarm(colour="turquoise3",cex = 1, size=2, alpha=0.6,
dodge.width=0.5 )+
  stat_compare_means(method="t.test")+
```

```
  stat_n_text()+
  theme_bw()+
  ylab("Male Growth Rate (mg/day)")+
  theme(axis.text.x=element_text(size=9))+
  stat_summary(fun=mean, colour="black", geom="errorbar", aes(ymax =
..y.., ymin = ..y..),width=0.75,linetype="dashed")+
  stat_summary(fun=mean, geom="text", vjust=-0.7,size=3,
aes(label=paste("Mean:",round(..y.., digits=1))))

female_only<-subset(Data_master, sex=="Female")
ggplot(data=subset(female_only, treatment %in% c("20~30","23~33"),
                   body.adult.mass.g!="" | !(is.na(body.adult.mass.g))),
       aes(x = treatment, y = as.numeric(body.adult.mass.g)*1000/
(as.numeric(pupation.eclosion.time)/24))) +
  geom_boxplot(col=c("dodgerblue","orange"))+
  geom_beeswarm(colour="palevioletred1",cex = 1, size=2, alpha=0.6,
dodge.width=0.5 )+
  stat_compare_means(method="t.test")+
  stat_n_text()+
  theme_bw()+
  ylab("Female Growth Rate (mg/day)")+
  theme(axis.text.x=element_text(size=9))+
  stat_summary(fun=mean, colour="black", geom="errorbar", aes(ymax =
..y.., ymin = ..y..),width=0.75,linetype="dashed")+
  stat_summary(fun=mean, geom="text", vjust=-0.7,size=3,
aes(label=paste("Mean:",round(..y.., digits=1))))


#Wing size against body mass
ggplot(data=subset(Data_master, treatment %in% c("20~30","23~33"),
                   body.adult.mass.g!="" | !(is.na(body.adult.mass.g))),
       aes(x=(body.adult.mass.g)*1000, y=area.average,
group=treatment,colour=treatment,fill=treatment))+
  #geom_abline(slope=1, intercept = 201, alpha=0.25, size=1)+
  scale_color_manual(values= c("#56B4E9","gold2"))+
  scale_fill_manual(values= c("#56B4E9","gold2"))+
  geom_point(alpha=0.5,size=2, aes(shape=sex))+
  geom_smooth(method=lm,alpha=0.2)+
  theme_bw()+
  #stat_cor(method = "pearson", label.x=5)+
  ylab("Wing Size (mm^2)")+
  xlab("Adult Body Mass (mg)")+
  theme(axis.text.x=element_text(size=14),
        axis.text.y=element_text(size=14),
        axis.title=element_text(size=16))+
  stat_cor(method = "pearson", label.x=40)+
  theme(panel.border = element_blank(),panel.grid.major = element_blank(),
        panel.grid.minor = element_blank(), axis.line =
element_line(colour="black"))


#linear models
lm_data= subset(Data_master,treatment %in% c("20~30", "23~33", "26~36"))
lm1=lm(formula = area.average ~ treatment +sex, data=lm_data)
summary(lm1)

lm2 = lm(formula= (body.adult.mass.g / area.average) ~ treatment + sex,
data=lm_data)
```

```
summary (lm2)
```
